# Hippocampal CA2 neurons disproportionately express AAV-delivered genetic cargo

**DOI:** 10.1101/2024.11.27.625768

**Authors:** Georgia M. Alexander, Bo He, Austin Leikvoll, Stephanie Jones, Rob Wine, Prakash Kara, Negin Martin, Serena M. Dudek

## Abstract

Hippocampal area CA2 is unique in many ways, largely based on the complement of genes expressed there. We and others have observed that CA2 neurons exhibit a uniquely robust tropism for adeno-associated viruses (AAVs) of multiple serotypes and variants. In this study, we aimed to systematically investigate the propensity for AAV tropism toward CA2 across a wide range of AAV serotypes and variants, injected either intrahippocampally or systemically, including AAV1, 2, 5, 6, 8, 9, DJ, PHP.B, PHP.eB, and CAP-B10. We found that most serotypes and variants produced disproportionally high expression of AAV-delivered genetic material in hippocampal area CA2, although two serotypes (AAV6 and DJ) did not. In an effort to understand the mechanism(s) behind this observation, we considered perineuronal nets (PNNs) that ensheathe CA2 pyramidal cells and, among other functions, buffer diffusion of ions and molecules. We hypothesized that PNNs might attract AAV particles and maintain them in close proximity to CA2 neurons, thereby increasing exposure to AAV particles. However, genetic deletion of PNNs from CA2 had no effect on AAV transduction. Next, we next considered the AAV binding factors and receptors known to contribute to AAV transduction. We found that the AAV receptor (AAVR), which is critical to transduction, is abundantly expressed in CA2, and knockout of AAVR nearly abolished expression of AAV-delivered material by all serotypes tested. Additionally, we found CA2 enrichment of several cell-surface glycan receptors that AAV particles attach to before interacting with AAVR, including heparan sulfate proteoglycans, N-linked sialic acid and N-linked galactose. Indeed, CA2 showed the highest expression of AAVR and the investigated glycan receptors within the hippocampus. We conclude that CA2 neurons are endowed with multiple factors that make it highly susceptible to AAV transduction, particularly to the systemically available PHP variants, including CAP-B10. Given the curved structure of hippocampus and the relatively small size of CA2, systemic delivery of engineered PHP or CAP variants could all but eliminate the need for intrahippocampal AAV injections, particularly when injecting recombinase-dependent AAVs into animals that express recombinases in CA2.

## INTRODUCTION

Hippocampal area CA2 is a distinct member of the hippocampal subfields, owing to a robust and selectively expressed set of genes (Lein et al., 2005; Dudek et al., 2016; Farris et al., 2019), an intriguing and unique synaptic plasticity profile (Zhao et al., 2007; Chevaleyre and Siegelbaum, 2010), and its role in social memory (Hitti and Siegelbaum, 2014; Alexander et al., 2016; Smith et al., 2016), social aggression (Leroy et al., 2018), and defensive behavior (Radzicki et al., 2024). We have observed in our own studies and in the literature another unique feature of CA2 neurons: the propensity of CA2 neurons to richly express genetic cargo carried by adenoassociated viruses (AAVs), even when AAVs are injected away from CA2 intrahippocampally and when injected systemically (Divoudi and Foster, 2019; Chan et al., 2017; Besnard et al., 2020; Simonnet at el., 2023, for example). This observation led us to ask why CA2 neurons preferentially express AAV-delivered genetic cargo to understand the nuances of CA2, capitalize on the use of AAVs in CA2, and possibly minimize off-target expression of AAVs intended for delivery to areas adjacent to, but not including, CA2. As AAVs are a heavily used tool for studying brain structure and function (see Nectow and Nestler, 2020, for review), identifying the mechanisms supporting spread and tropism of AAVs can help guide selection of AAV serotypes and variants.

AAVs are single-stranded DNA viral particles that gain entry to cells through interactions of capsid proteins with cell surface molecules, including glycans and proteinaceous receptors, leading to internalization and subsequent trafficking to the nucleus and expression of genetic material (Dhungel et al., 2020). The various serotypes and variants of AAVs have a distinct complement of capsid domains that are thought to preferentially interact with distinct primary receptors and co-receptors to facilitate cell internalization (Meyer and Chapman, 2022). In particular, the AAV receptor (AAVR), encoded by the *KIAA0319L* gene, has been shown to be required for AAV transduction in cell lines by several serotypes and *in vivo* by the AAV2 serotype (Pillay et al., 2016). Because we observe prominent AAV tropism with various serotypes and variants in CA2, we hypothesized that CA2 neurons may richly express factors that facilitate AAV transduction and/or the AAVR itself. However, because different serotypes and variants have different cell-surface interaction domains on their capsids (Pillay et al., 2017), we hypothesized that CA2 neurons may be selectively vulnerable to AAV transduction in a serotype/variant specific manner. Therefore, we examined tropism of a wide range of AAV serotypes/variants following intrahippocampal or systemic administration.

CA2 neurons are also unique in that they are surrounded by a specialized extracellular matrix called perineuronal nets (PNNs). Indeed, CA2 neurons are among the few glutamatergic cell types in the brain to possess PNNs, and the only glutamatergic cells among the hippocampal CA fields with them (Brückner et al., 2003; Costa et al., 2007; Carstens et al., 2016). Otherwise, PNNs are found surrounding inhibitory parvalbumin-expressing interneurons in hippocampus and throughout the CNS (Härtig et al., 1992; Celio, 1993). PNNs are composed of negatively charged chondroitin sulfate proteoglycan (CSPG) molecules bound together by link proteins and glycoproteins, tethered by hyaluronic acid molecules and attached to the cell membrane by hyaluronan synthase (Sanchez et al., 2023). Among the functions of PNNs, they are thought to buffer the diffusion of molecules and ions at synapses and hold synaptic components in place around the cell soma and proximal dendrites in their spongy lattice-like structure (Morawski et al., 2015; Fawcett et al., 2022). Given these structural and functional attributes, we hypothesized that PNNs surrounding CA2 neurons may capture and retain AAV particles traveling in the extracellular space, resulting in greater exposure of CA2 neurons to AAV particles, and, in turn, greater AAV transduction. Additionally, AAV particles carry positively charged moieties on their capsid surface, which may interact with the negatively charged CSPGs. Similar extracellular matrix molecule to CSPGs with a similarly negative charge, heparan sulfate proteoglycan (HSPGs), are known to interact with AAVs to facilitate internalization (Opie at al., 2003), further supporting the notion that CSPGs of PNNs may contribute to our observation that CA2 neurons are disproportionately transduced and, thus, disproportionately express AAV cargos.

In this study we asked whether CA2 neurons differ in their AAV tropism in a serotype/variant specific manner and what mechanism may account for the preferential expression. Through intrahippocampal and retro-orbital injection of AAVs of several serotypes and variants, we found that most of the tested serotypes/variants disproportionately transduce CA2, although some do not. In addition, using a conditional knock-out mouse strain for the primary CSPG in CA2 neurons to eliminate PNNs selectively from CA2, we show that PNNs do not contribute to the preferential expression of AAV material by CA2 neurons. Instead, we found the CA2 neurons are endowed with several other proteins and molecules that facilitate AAV transduction, including the AAV receptor (AAVR), primary glycan receptors, and the AAV2 co-receptor, fibroblast growth factor receptor 1 (FGFR1). One serotype, AAV6, seems capable of robust transduction of neurons with low expression levels of factors found in CA2 neurons and may also provide a useful tool for avoiding unintended expression in CA2. Finally, systemically available AAVs, including the PHP variants, and especially CAP-B10, provide robust and selective transduction of CA2 neurons, supporting the use of these variants for non-invasive AAV administration in CA2 research.

## METHODS

### Animals and injections

Adult mice (8-12 weeks at the time of injections) of the following strains were used: C57BL/6J (Jackson Laboratory; strain #000664), Amigo2iCreERT2; ROSA-tdTomato (Alexander et al., 2018; ROSA-tdTomato: Jackson Laboratory strain #007909), AAVR KO (MMRRC; stock # 043535-UCD), EMX Cre+; *Nr3C2* fl/fl (EMX Cre; MR fl/fl; McCann et al., 2021; McCurley et al., 2012; EMX Cre: Jackson Laboratory; strain #005628), Amigo2 Cre; *Acan* fl/fl (Alexander et al., 2024). Amigo2iCreERT2; ROSA-tdTomato were treated with tamoxifen (100 mg/kg IP, daily for 3 days) to permit Cre activity and thus express tdTomato fluorophore in CreER expressing cells. CAP-B10-hSyn-jGCaMP8s was injected into 2 C57BL/6J mice at the age of 4 weeks. Mice were housed under a 12:12 light/dark cycle with access to food and water *ad libitum*. Mice were group housed and were naïve to any treatment, procedure, or testing when beginning the experiments described here. All procedures were approved by the NIEHS and University of Minnesota Animal Care and Use Committees and were in accordance with the National Institutes of Health guidelines for care and use of animals. Both males and females, as defined by the presence or absence of the Y chromosome, were used in these experiments. A total of 29 C57BL/6J (28 male, 1 female), 5 male Amigo2iCreERT2; ROSA-tdTomato, 28 AAVR KO (15 male, 13 female), 32 EMX Cre; MR fl/fl (16 male, 16 female) and 24 Amigo2 Cre; *Acan* fl/fl (14 male, 10 female) animals were used. For the AAVR KO, EMX Cre; MR fl/fl and Amigo2 Cre; *Acan* fl/fl experiments, which included males and females, we found no differences in our experimental measures according to sex, so data from males and females were combined. Experimenters were blinded to genotype at the time of experiments and during data analysis.

CAP-B10-hSyn-jGCaMP8s was generated by the University of Minnesota viral vector and cloning core using plasmids from Addgene (CAP-B10: #175004; hSyn-jGaMP8s: #162374). All other viruses were generated by the NIEHS viral vector core laboratory using plasmids from Addgene (PHP.B: #103002; PHP.eB: #103005; hSyn-EGFP: #50465) or Cell Biolabs Inc. (AAV1: #VPK-421; AAV2: #VPK-422; AAV5: # VPK-425; AAV6: #VPK-426; AAV8: #VPK-428). AAV9 was a generous gift of UPenn Vector Core (pAAV2/9). AAVDJ was a generous gift of the laboratory of Mark Kay, Stanford University (pAAV2/DJ). For retro-orbital sinus injections, animals were anesthetized with isoflurane, injected retro-orbitally with a 30-ga needle, and returned to the home cage on heat until completely recovered from anesthesia. For the CAP-B10 serotype, a total of 4×10^11^ GC was injected. For the PHP.B and PHP.eB serotypes, a total of 1×10^11^ GC was injected. For intrahippocampal virus injection surgery, animals were anesthetized with ketamine (100 mg/kg, IP) and xylazine (7 mg/kg, IP), then positioned in a stereotaxic apparatus. An incision was made in the scalp, a hole was drilled over the target region for AAV infusion, and a 30-ga cannula connected to a Hamilton syringe by a length of tube was lowered into hippocampus. For injections targeting all areas of hippocampus for assessment of expression level in each subfield (i.e., Fig. 1), injection coordinates were, in mm: −2.3 AP, ±2.3 ML, −1.8 DV relative to bregma (Paxinos and Franklin, 2001). Injections were made bilaterally, and each hemisphere represents an observation (N). Within a given animal, the same serotype/variant of AAV was injected on both the right and left hippocampus. For injections with AAV6-hSyn-GFP targeting CA2 specifically (i.e., Fig. 7), injection coordinates were, in mm: −1.8 AP, ±2.05 ML, −1.8 DV. For all intrahippocampal injections, AAV titers were 1×10^12^ GC/ml, and 250 nl of AAV was injected at an injection rate of 50 nl/min. Following injection, the scalp was sutured shut and animals returned to a clean cage on heat until mobile and alert. Animals were administered buprenorphine (0.1 mg/kg, SQ) for pain.

**Figure 1.**
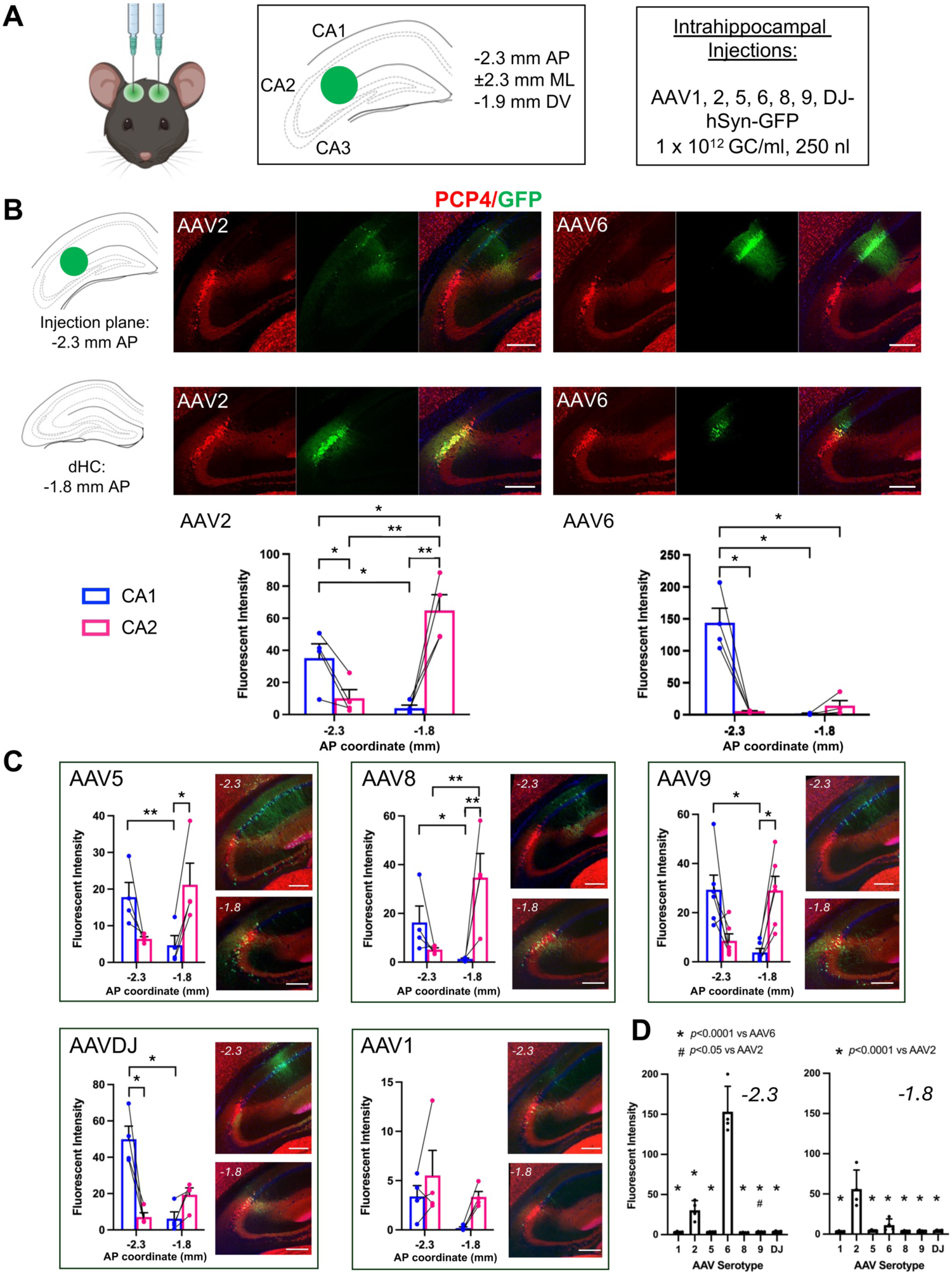
Intrahippocampally injected AAVs showed differential tropism for CA2 according to serotype or variant. A. Male C57BL/6J mice were injected bilaterally with AAV-hSyn-GFP of various serotypes/variants at the listed coordinates, all relative to Bregma. The specific serotypes and titer, held constant for all serotypes, are listed, and the volume injected in each instance was 250 nl. B. Representative images of GFP delivered by AAV2 (left) or AAV6 (right) serotypes at each of the injection site place (top row, −2.3 mm AP) and a more anterior plane (bottom row, −1.8 mm AP), defined as dorsal hippocampus (dHC) here. Plotted below are mean fluorescence intensities of GFP in the CA2 pyramidal cell layer, defined by PCP4 expression, and a similar area of CA1 pyramidal cell layer at each AP plane. AAV2 and AAV6, which showed the most robust expression of the serotypes, displayed opposite affinity for CA2 expression, with AAV2 showing the greatest GFP expression in CA2 of dHC and AAV6 showing the greatest GFP expression at the injection site in CA1. C. Mean fluorescence intensities for GFP delivered by the remaining AAVs and representative images from each plane of section. For measurements shown in B and C, image acquisition settings were held constant for all samples within a serotype, but settings differed across serotypes due to the relative expression level afforded by each. D. Samples were re-imaged using identical image acquisition settings for all samples across all serotypes to demonstrate the relative level of GFP expression with each AAV. The area of maximum fluorescence intensity, regardless of hippocampal subfield, was collected at each AP coordinate plane listed and compared across serotype (−2.3 AP: F(6,23)=84.91, *p*<0.0001; −1.8 AP: F(6,23)=18.79, *p*<0.0001, one-way ANOVAs with Tukey’s multiple comparisons tests). Results of repeated measures two-way ANOVAs associated with B and C are shown in Table 1. Results of Sidak’s multiple comparisons tests following two-way ANOVAs (B-C) and Tukey’s tests following one-way ANOVAs (D) shown on graphs. Replicates reflect brain hemispheres (two per animal). \**p*<0.05, \*\**p*<0.01. All scale bars = 250 μm.

**Table 1.** Statistics Associated with Figure 1.

| **Serotype** | **Subfield** | **AP plane** | **Interaction** |
| --- | --- | --- | --- |
| AAV2 | F(1,3)=3.92, p=0.14 | F(1,3)=1.71, p=0.28 | F(1,3)=266.9, p=0.0005 *** |
| AAV6 | F(1,3)=20.92, p=0.019 * | F(1,3)=21.54, p=0.019 * | F(1,3)=62.17, p=0.0043 ** |
| AAV5 | F(1,3)=3.78, p=0.15 | F(1,3)=0.10, p=0.77 | F(1,3)=14.74, p=0.031 * |
| AAV8 | F(1,3)=1.76, p=0.28 | F(1,3)=0.82, p=0.43 | F(1,3)=106.9, p=0.0019 ** |
| AAV9 | F(1,5)=0.33, p=0.59 | F(1,5)=0.78, p=0.42 | F(1,5)=22.05, p=0.0054 ** |
| AAVDJ | F(1,3)=9.71, p=0.053 | F(1,3)=7.22, p=0.075 | F(1,5)32.92, p=0.011 * |
| AAV1 | F(1,3)=0.42, p=0.56 | F(1,3)=3.78, p=0.15 | F(1,3)=5.54, p=0.10 |
| Repeated-measures two-way ANOVAs |  |  |  |

### Tissue processing and imaging

Three weeks after AAV injection (or 5-8 weeks following CAP-B10-hSyn-jGCaMP8s injection), animals were euthanized with Fatal Plus (sodium pentobarbital, 50 mg/mL; >100 mg/kg) and perfused transcardially with phosphate buffered saline (PBS) followed by 4% paraformaldehyde. Brains were post-fixed in 4% paraformaldehyde overnight before being transferred to PBS. Brains were sectioned on a vibratome at 40 μm and stored in PBS with 0.1% sodium azide.

For immunohistochemistry, sections were rinsed in PBS, boiled in deionized water (for reactions including AAVR only), further rinsed in PBS then PBS with 0.1% triton X-100 (PBST), blocked for 1 hr with 5% normal goat serum in PBST, then incubated in blocking solution plus primary antibodies or lectins overnight at 4°C. Primary antibodies included PCP4 (Invitrogen, PA5-52209, 1:500, raised in rabbit, and Synaptic Systems, 480 004, 1:500, raised in Guinea pig), GFP (Invitrogen, A10262, 1:1000, raised in chicken), AAVR (Novus, NBP2-57263, 1:500, raised in rabbit), ZnT3 (Synaptic Systems, 197 004, 1:500, raised in Guinea pig), aggrecan (Millipore, AB1031, 1:500, raised in rabbit), brevican (Invitrogen, MA5-27639, 1:500, raised in mouse), neurocan (Invitrogen, PA5-47779, 1:500, raised in sheep), versican (Invitrogen, MA5-27638, 1:500, raised in mouse), FGFR1 (Invitrogen, PA5-86246, 1:1000, raised in rabbit), LY6A (Invitrogen, 14-5981-82, 1:500, raised in rat), WFS1 (Proteintech, 11558-1-AP, 1:500, raised in rabbit), and 10E4 heparan sulfate (amsbio, 370255-B-50-H, 1:200, biotinylated). Lectins included *Wisteria floribunda* agglutinin (WFA; Vector labs, B-1355, 1:1000, biotinylated), *Sambucus nigra* agglutinin (SNA; Vector labs, B-1305-2, 1:1000, biotinylated), and *Maackia amurensis* lectin I (MAL I; Vector Labs, B-1315, 1:500). The next day, brains were washed in PBST and incubated in blocking buffer plus secondary antibodies, all raised in goat (except anti-sheep), purchased from Invitrogen and using a dilution of 1:500, including anti-chicken 488 (A11039), anti-rabbit 488 (A11034), anti-rabbit 568 (A11011), anti-mouse 488 (A32723), anti-Guinea pig 568 (A11075), anti-Guinea pig 633 (A21105), donkey anti-sheep 488 (Invitrogen, A11015), anti-rat 568 (A11077). Streptavidin conjugates included 488 (Invitrogen, S32354) or 568 (Invitrogen, S11226). Sections were then washed and mounted with mounting medium containing DAPI (Vector labs, H-1500). Brains were cleared for 3D imaging using the Fast 3D Clear protocol (Kosmidis et al., 2021). Briefly, brains were washed in PBS then deionized H_2_O, dehydrated and delipidated in a series of tetrahydrofuran dilutions then cleared brains in an aqueous iohexol-based solution at 37°C. Cleared brains were maintained at 4°C in clearing solution protected from light until imaging.

Images of brain sections were acquired on Zeiss LSM 880 and Zeiss 980 confocal microscopes. Cleared brains were imaged on a Zeiss Lightsheet 7, with brains immersed Cargille oil. Within experiments comparing fluorescence intensity across genotype or brain areas, image acquisition settings were held constant for all samples, unless otherwise stated.

### Data Analysis

Fiji software (Schindelin et al., 2012) was used to quantify mean fluorescence intensity from acquired images of stained brain section. For expression of intrahippocampally injected AAVs (i.e., Fig. 1, Supp. Fig. 1), measurements were taken from brain sections corresponding to mouse brain atlas planes −2.3 mm, and −1.8 mm AP (Paxinos and Franklin, 2001). Regions of interest (ROIs) were created in the pyramidal cells layer of each subfield, using a similar area for each ROI (area of CA2 ±5%). Specifically, CA2 ROIs were generated using PCP4 expression in the pyramidal cell layer. The area of CA2 ±5% dictated the size of the ROI in each of the remaining subfield cell body layers, and ROIs were positioned over the area of greatest GFP expression in that subfield. For all other quantifications, measurements were taken from 2-3 hippocampi per animal, with ROIs created in the pyramidal cell layers and values averaged for an N. Area CA2 was defined by PCP4 expression for all measurements, except those in MR fl/fl animals, for which the pyramidal cells at the distal end of the ZnT3-positive mossy fibers (200 µm in length ROI) was used to define CA2 because no known CA2 markers can be detected (McCann et al., 2021). All mean fluorescence values were background-subtracted. Fiji software was used for image processing of cleared brain image files, and Imaris software (Oxford instruments) was used to generate videos of cleared brain.

Graphpad Prism 10 (Graphpad) was used for all statistical analyses. Two-factor grouped comparisons were analyzed using repeated measured (RM) two-way ANOVAs, and post-hoc test *p* value results were corrected for multiple comparisons. Single factor comparisons with two groups were analyzed using Student’s t-tests. Equality of variance was tested by F-tests, and normality was tested using D’Agostino and Pearson tests. Specific statistical tests used and the results of the tests are listed in the figure legends. Significance was set at *p*<0.05. For all graphs, mean and standard errors (SEM) are plotted, and dots in graphs represent individual animals/observations.

## RESULTS

### Most AAV serotypes and variants preferentially target dorsal CA2 neurons

To address whether CA2 neurons show ubiquitous or selective preference for AAV transduction, we surveyed several serotypes and variants following either local intrahippocampal injection or systemic retro-orbital injection. All AAVs carried a genetic payload of hSyn-GFP, except for one AAV, which carried hSyn-jGCaMP8s. Of the 10 serotypes and variants that we tested following local or systemic injection, 7 showed preferential transduction of dorsal CA2 neurons within hippocampus.

For AAVs injected intrahippocampally, titers, injection volumes and injection flow rates were held constant. We selected an injection site equidistant from each subfield, including CA1, CA2, CA3 and DG, which required penetration of the CA1 pyramidal cell layer (Fig. 1A). Brains were collected three weeks after injections. Upon examination of the brain sections, we found two populations of cells that predominantly expressed GFP: in CA1 near the site of cannula penetration and in CA2 at more anterior planes. To characterize and quantify our observations, we measured the GFP fluorescence intensity in each subfield at two coronal planes of section: the injection plane (−2.3 mm AP) and 0.5 mm anterior to that plane (−1.8 mm AP).

Two serotypes, AAV2 and AAV6, were striking in their robustness of expression and their opposing affinity for expression location (Fig. 1B, Table 1, Supp. Fig. 1). At the injection plane, AAV2 transduced CA1 neurons to a greater extent than other subfields. However, at the more anterior plane, AAV2-delivered GFP expression was greater in CA2 than any other subfield and was also greater than that in CA1 at the injection plane. Thus, AAV2 shows a significant affinity for dorsal CA2 neurons. Conversely, AAV6-delivered GFP showed the greatest expression level in CA1 at the plane of injection, and minimal GFP expression was seen in dorsal CA2 at the more anterior plane. Thus, AAV6 shows a significant affinity for CA1, or at least true to the site of cannula penetration and injection.

Of the remaining serotypes/variants injected into hippocampus, AAV5, AAV8 and AAV9 showed an expression pattern similar to that of AAV2, with a substantial (though not significant) level of expression in CA1 at the injection plane but a significant preference for CA2 expression at the more anterior plane (Fig. 1C, Table 1). By contrast, AAVDJ resembled AAV6 in its relative selectivity for targeted expression, with a significant preference for expression in CA1 at the injection plane. AAV1 was difficult to characterize due to the low expression levels that it permitted.

The overall expression levels of GFP differed according to serotype/variant. Therefore, in the results described above, image acquisition settings were held constant within a serotype/variant but differed across them. To compare the relative GFP expression levels after injection with each of the serotypes/variants, we re-imaged the brain sections with acquisition settings held constant across samples and serotypes/variants. Specifically, those setting used for AAV6 were used for all samples because it produced the greatest fluorescence signal. In doing so, we found that expression level of AAV6 at the plane of injection (−2.3 mm AP) was significantly greater than all other serotypes/variants.

However, at the more anterior dorsal hippocampus plane, GFP expression level using AAV2 was significantly greater than all other serotypes/variants (Fig. 1D). These findings support the conclusion that CA2 neurons have a high affinity for many, but not all, serotypes and variants of AAV. On a more qualitative level, we also observed a differential degree of spread for the various AAVs. For example, AAV6-delivered GFP expression is largely confined to a small area, whereas AAV5 produced a large spread of GFP-expressing neurons.

To eliminate the variable of intrahippocampal injection site and also to examine CA2 expression of further AAV variants, we employed the PHP.B, PHP.eB and CAP-B10 AAV variants, which can be administered retro-orbitally and can pass the blood-brain barrier (Chan et al., 2017). The PHP.B and PHP.eB variants carried hSyn-GFP and CAP-B10 carried hSyn-jGCaMP8s. As with several of the intrahippocampally-injected AAVs, we found that each of the retro-orbitally injected AAVs produced robust CA2 expression of GFP (Fig. 2, Supp. Video 1). For PHP.B and PHP.eB, expression of GFP was significantly greater in CA2 than in any other subfield (Fig. 2A-B), but GFP was also richly expressed in cortical and sub-cortical areas (although not quantified here). CAP-B10 appeared to show even greater selectivity for CA2 within hippocampus (Fig. 2C, Supp. Fig. 2), and cortical and subcortical expression was present, although less so than with the PHP variants (also not quantified here).

**Figure 2.**
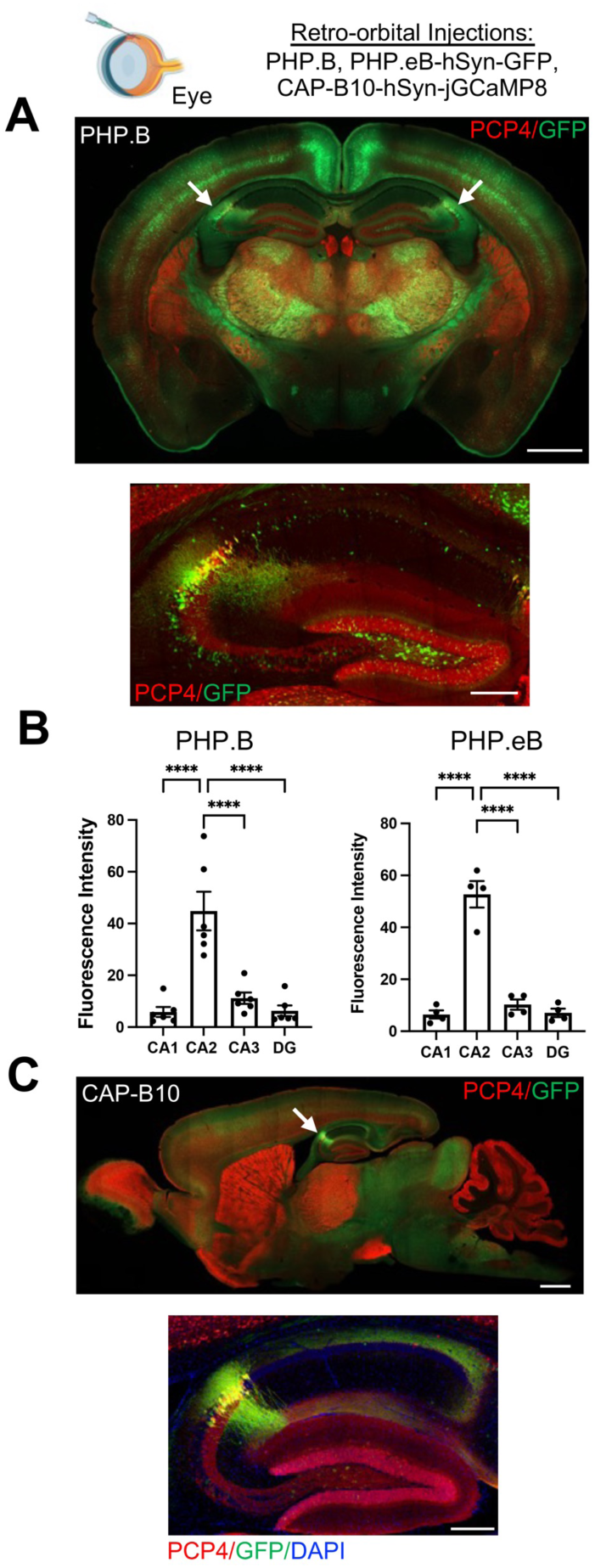
PHP.B, PHP.eB, and CAP-B10 AAV variants, administered by retro-orbital injection, preferentially target CA2 neurons. C57BL/6J mice were injected retro-orbitally with hSyn-GFP packaged in either PHP.B or PHP.eB serotypes (A-B) or hSyn-jGCaMP8s packaged in CAP-B10 (C). A total of 1 x 10^11^ GC of PBP.B or PHP.eB or 4 x 10^11^ GC CAP-B10 was administered, with volume adjusted based on the titer. The representative images show a whole-brain coronal (A) or sagittal (C) sections immunostained for the CA2 marker, PCP4, and GFP following injection with PHP.B (A) or CAP-B10 (C) variants. CA2 is marked by the white arrows. Below each whole-brain image, an image of hippocampus only is shown. B. GFP fluorescence intensity was quantified in each hippocampal subfield for PHP.B and PHP.eB-injected animals. GFP expression in CA2 was significantly greater than expression in any other hippocampal subfield for both serotypes (PHP.B: F(3,15)=37.35, *p*<0.0001; PHP.eB: F(3,9)=107.8, *p*<0.0001). Results of Tukey’s multiple comparisons tests shown on graphs. Scale bars = 1 mm for whole brain images and 250 μm for hippocampal images.

### AAVR is required for effective AAV transduction

In attempts to identify why CA2 neurons preferentially express AAV-delivered genetic material, we considered the receptors and factors that contribute to AAV transduction. We began with the AAV receptor (AAVR; Pillay et al., 2016; gene name *KIAA0319L*). We immunostained tissue from C57BL/6J mice for AAVR and found that expression was significantly higher in CA2 than surrounding subfields. In addition, cells with greater AAVR expression also appeared to express greater GFP expression following PHP.B-hSyn-GFP injection (Fig. 3A-B). We then asked whether AAVR is required for CA2 expression of AAV-delivered genetic material by employing AAVR KO animals, testing systemically available and intrahippocampally-injected serotypes/variants. First, we retro-orbitally delivered PHP.B or PHP.eB-hSyn-GFP to AAVR KOs and WT littermates, and we found that GFP expression was abolished in AAVR KO animals. GFP expression level was significantly less in AAVR KO animals than WTs in all hippocampal subfields, and indeed throughout the brain (Fig. 3C-D). Next, we examined GFP fluorescence in WT and AAVR KO animals following intrahippocampal injection of the previously investigated serotypes/variants (shown in Fig. 1). As with the intrahippocampal injections in C56BL/6J mice, image acquisition settings varied according to serotype/variant but were held constant within each serotype/variant. We found that for each of the serotypes/variants, GFP expression was significantly less (greater than 90% less) in AAVR KO animals than in WTs (Fig. 3E*i*), indicating a substantial role, if not a prerequisite, of AAVR in AAV transduction of all serotypes/variants tested here. However, we again noted that AAV6 produced a considerable GFP signal, albeit significantly less in the AAVR KO than the WT (Fig 3E*ii*). To explore this further, we reimaged only the AAVR KO tissue using identical image acquisition settings for all samples and measured the GFP fluorescence intensity. We found that AAV6 produced significantly greater GFP expression in AAVR KOs than in any of the other intra-hippocampally injected serotypes/variants (Fig. 3E*iii*), indicating that although AAVR is required for the full extent of AAV6’s robustness, even in its complete absence, AAV6 can transduce neurons considerably well. Consistent with this idea, AAV6 produced the largest overall GFP signal in CA1 of C57BL/6J mice (Fig. 1A), yet AAVR expression in CA1 is on the low end of that seen in hippocampus (Fig 3A).

**Figure 3.**
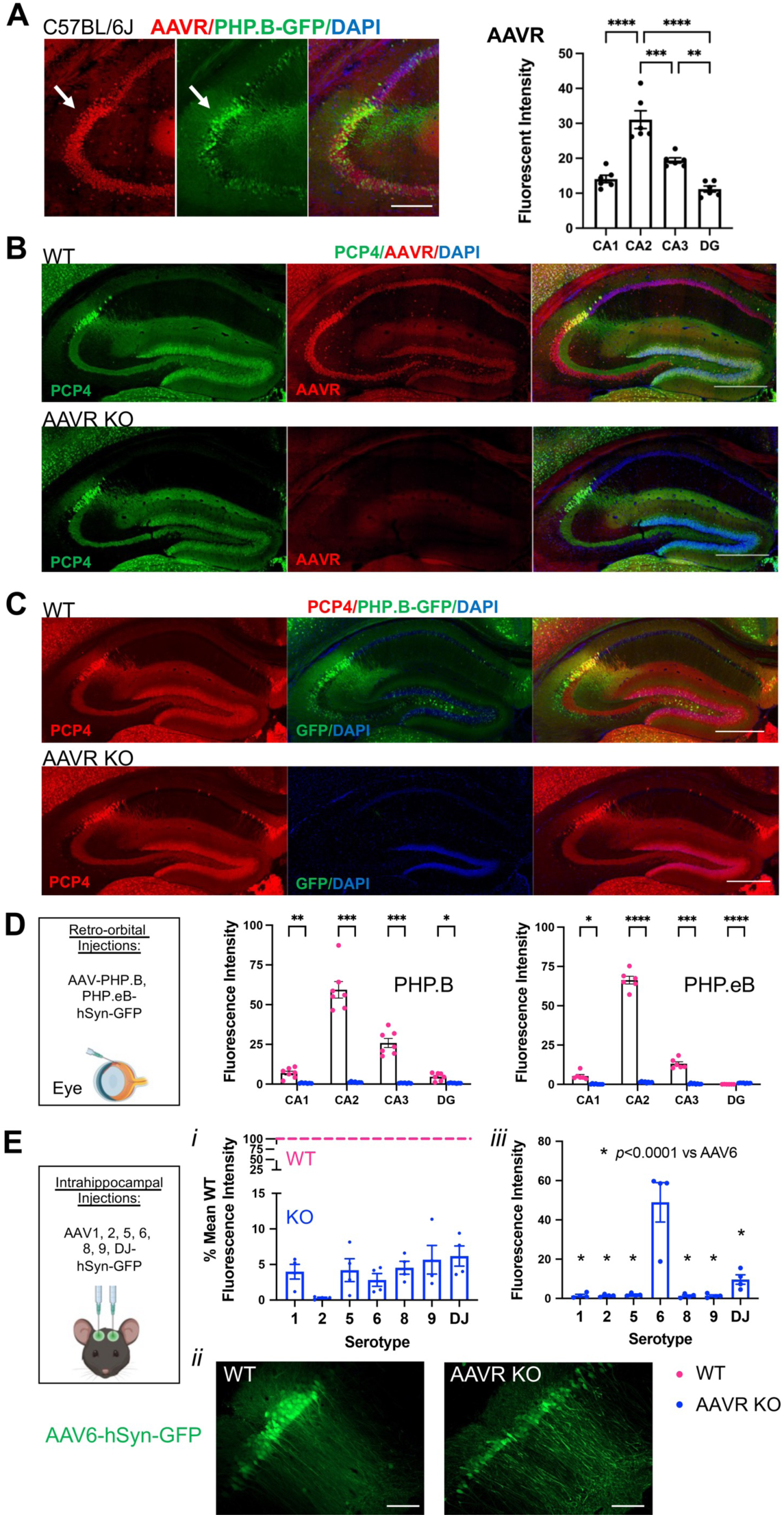
CA2 highly expresses AAVR, and AAVR KO animals lack AAV expression. A. Immunofluorescence for AAVR and GFP in PHP.B-hSyn-GFP-injected animals shows overlap of high AAVR expression and GFP expression in CA2 (arrows). Quantification of AAVR fluorescence intensity shows significantly greater AAVR expression in CA2 than surrounding subfield (F(3,20)=34.80, *p*<0.0001; one-way ANOVA, results of Tukey’s post hoc tests shown on graph). B-C. Whereas WT animals show robust AAVR (B) and AAV-delivered GFP expression (C) in CA2, AAVR KO animals lack both AAVR (B) and AAV-delivered GFP expression (C). D. Quantification of GFP fluorescence intensity in WT and AAVR KO animals following retro-orbital injection of PHP.B or PHP.eB-hSyn-GFP. GFP expression was significantly lower in all subfields in the AAVR KO animals compared with WT littermates (PHP.B: genotype: F(1,12)=107.2, *p*<0.0001, subfield: F(1.4,16.5)=118.7, *p*<0.0001, interaction: F(3,36)=114.1, *p*<0.0001; PHP.eB: genotype: F(1,12)=1399, *p*<0.0001, subfield: F(1.4,16.9)=561.2, *p*<0.0001, interaction: F(3,36)=530.1, *p*<0.0001, two-way ANOVAs with Geisser-Greenhouse correction, results of Bonferroni multiple comparisons tests shown on graphs). Tissue from PHP.B and PHP.eB injections were processed and analyzed independently, so data cannot be directly compared between these two serotypes. E. Quantification of GFP fluorescence following intrahippocampal injection of AAV1, 2, 5, 6, 8, 9 or DJ-hSyn-GFP. Coordinates for injections: −2.3 AP, +/- 2.3 ML, −1.9 DV. E*i*. In comparing expression between genotypes, image acquisition settings were held constant within serotypes but differed across serotypes. Data are shown as the fluorescence intensity in KOs relative to mean expression in WTs for each serotype. Based on raw fluorescence intensity values, all serotypes had significantly lower expression in AAVR KOs than WTs (AAV1: t(6)=11.02, *p*<0.0001; AAV2: t(6)=2.91, *p*=0.026; AAV5: t(6)=8.06, *p*=0.0002; AAV6: t(6)=34.36, *p*<0.0001; AAV8: t(6)=4.20, *p*=0.0057, AAV9: t(6)=6.39, *p*=0.0007; AAVDJ: t(6)=2.46, *p*=0.049; two-tailed unpaired t-tests). E*ii*. GFP expression after delivery via AAV6 in AAVR WT and AAVR KO animals, although using separate microscope settings, meant to demonstrate the presence of GFP-expressing cells despite the lower overall expression level in the absence of the AAVR. E*iii*. Images were acquired a second time from KO animals only with image acquisition setting held constant across all serotypes, and GFP fluorescence intensity was quantified, to identify the most highly expressed AAV serotype despite the absence of AAVR. AAV6-delivered GFP showed the greatest fluorescence intensity of all intrahippocampally injected AAVs in AAVR KO animals (F(6,21)=20.38, *p*<0.0001, one-way ANOVA, results of Tukey’s multiple comparisons tests shown on graph). Scale bar in A=250 μm, B,C=500 μm, E=100 μm. *p<0.05, **p<0.01, ***p<0.001, ****p<0.0001.

From other studies in our lab in which we used conditional knockout mice for the mineralocorticoid receptor (MR; EMX Cre; MR fl/fl), we have anecdotally noted that CA2 neurons of EMX Cre+ MR fl/fl mice do not express AAV-delivered genetic material to the same extent as CA2 neurons in Cre-animals. In fact, we had found it difficult to infect CA2 neurons using the serotypes we commonly use (e.g., AAV2, AAV5). In this study, we took advantage of that anecdotal finding to determine what could be different about CA2 in the KO animals that hinders AAV transduction.

We first aimed to quantify the differential expression of GFP following AAV injection in the Cre- and Cre+ MR fl/fl animals, and to eliminate the variable of intrahippocampal injection site, we used retro-orbital injections of PHP.B-hSyn-GFP. Consistent with our anecdotal observations, we found that CA2 GFP expression was significantly less in Cre+ MR fl/fl animals than in Cre-controls (Fig. 4A-B). Interestingly, we also found that GFP expression was significantly greater in dentate granule cells of Cre+ than Cre-animals, although it is unclear why. We previously reported that expression of common protein markers of CA2 are absent in Cre+ MR fl/fl animals (McCann et al., 2021), which for the current study required us to use the anatomical localization of CA2 pyramidal cells at the distal end of the ZnT3-expressing mossy fibers to identify CA2 (Fig. 4A). Given the decreased GFP expression in CA2 of Cre+ MR fl/fl animals, we probed for the presence and expression level of AAVR and found that expression was significantly less in CA2 of Cre+ MR fl/fl animals than Cre-animals (Fig. 4D), again pointing to a significant role of AAVR in preferential tropism toward CA2.

**Figure 4.**
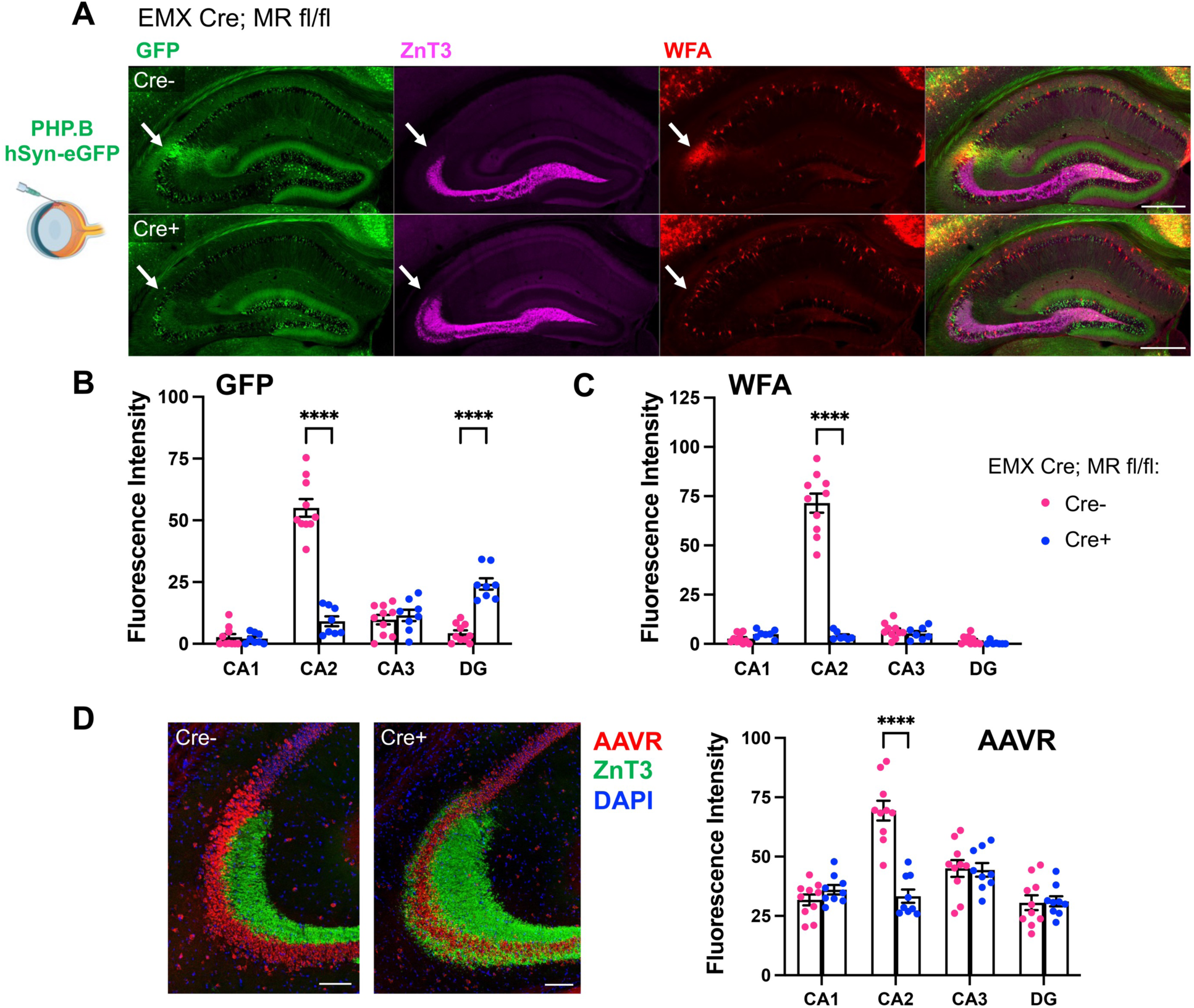
Mineralocorticoid receptor conditional KO animals (EMX Cre+; MR fl/fl) show less AAV-delivered GFP and less AAVR expression in CA2 than WT Cre-littermates. A. After retro-orbital injection with PHP.B-hSyn-GFP, Cre-animals showed robust expression in CA2 pyramidal cells (white arrows), found at the distal end of the mossy fibers, labeled by ZnT3. CA2 neurons also colocalize with perineuronal nets, as labeled by WFA. Cre+; MR fl/fl animals, which are known to lack CA2 perineuronal nets and other markers of CA2 pyramidal cells, showed significantly less GFP expression in CA2. As molecular markers of CA2 neurons are not expressed in CA2 of MR KO animals, we defined CA2 as the pyramidal cells at the distal end of the ZnT3-labeled mossy fibers (white arrows). B. Quantification of GFP fluorescence intensity in the cell body layer of each hippocampal subfield showed significantly decreased GFP expression in CA2 and significantly increased expression in DG of Cre+ MR fl/fl animals (subfield: F(3,48)=78.27, *p*<0.0001; genotype: F(1,16)12.20, *p*=0.0030; interaction: F(3,48)=98.02, *p*<0.0001). C. Quantification of WFA fluorescence intensity in the cell body layers. WFA was significantly decreased in CA2 of Cre+ MR fl/fl animals (subfield: F(3,45)=107.1, *p*<0.0001; genotype: F(1,15)=157.6, *p*<0.0001; interaction: F(3,45)=102.8, p<0.0001). D. Immunofluoresence for AAVR was significantly decreased in CA2 of Cre+ MR fl/fl animals compared with Cre-WTs (subfield: F(3,51)=75.29, *p*<0.0001; genotype: F(1,17)=4.35, *p*=0.052; interaction: F(3,51)=73.99, *p*<0.0001). Repeated-measured two-way ANOVAs with Bonferroni multiple comparisons used for all analyses, and results of multiple comparisons tests are shown on graphs. Scale bar in A=500 μm, D=100 μm.

### Perineuronal nets are not required for AAV transduction

One of the identifying markers of CA2 that is absent in Cre+ MR fl/fl animals caught our attention as potentially contributing to the propensity of CA2 neurons to express AAV-delivered material: Cre+ MR fl/fl animals lack aggrecan (Acan), the primary component of perineuronal nets (PNNs; McCann et al., 2021). Accordingly, Cre+ MR fl/fl animals lack PNNs, labeled by WFA, in CA2 (Fig. 4A, C). We considered the possibility that PNNs contribute to CA2’s propensity to express AAV-delivered material. To test this idea, we stained tissue from PHP.B-hSyn-GFP-injected C57BL/6J animals for WFA and found that in addition to the GFP-expressing cells in CA2 being surrounded by WFA, other cells outside of CA2 that stained for WFA also had GFP expression (Fig. 5A). Next, we asked whether animals lacking PNNs in CA2, but not other CA2 markers, would show decreased AAV-delivered GFP expression in CA2. For this, we used Amigo2 Cre; *Acan* fl/fl animals, in which *Acan* is deleted from CA2 neurons but not surrounding interneurons (Alexander et al., 2024; Supp. Fig. 2), and we again used retro-orbital injection of PHP.B-hSyn-GFP. We found that despite the absence of PNNs from CA2 pyramidal cells of Amigo2 Cre+; *Acan* fl/fl animals, GFP expression was not significantly decreased in CA2 (Fig. 5B-D, Supp. Video 2). Therefore, PNNs do not appear to contribute to the high affinity of CA2 neurons for AAVs.

**Figure 5.**
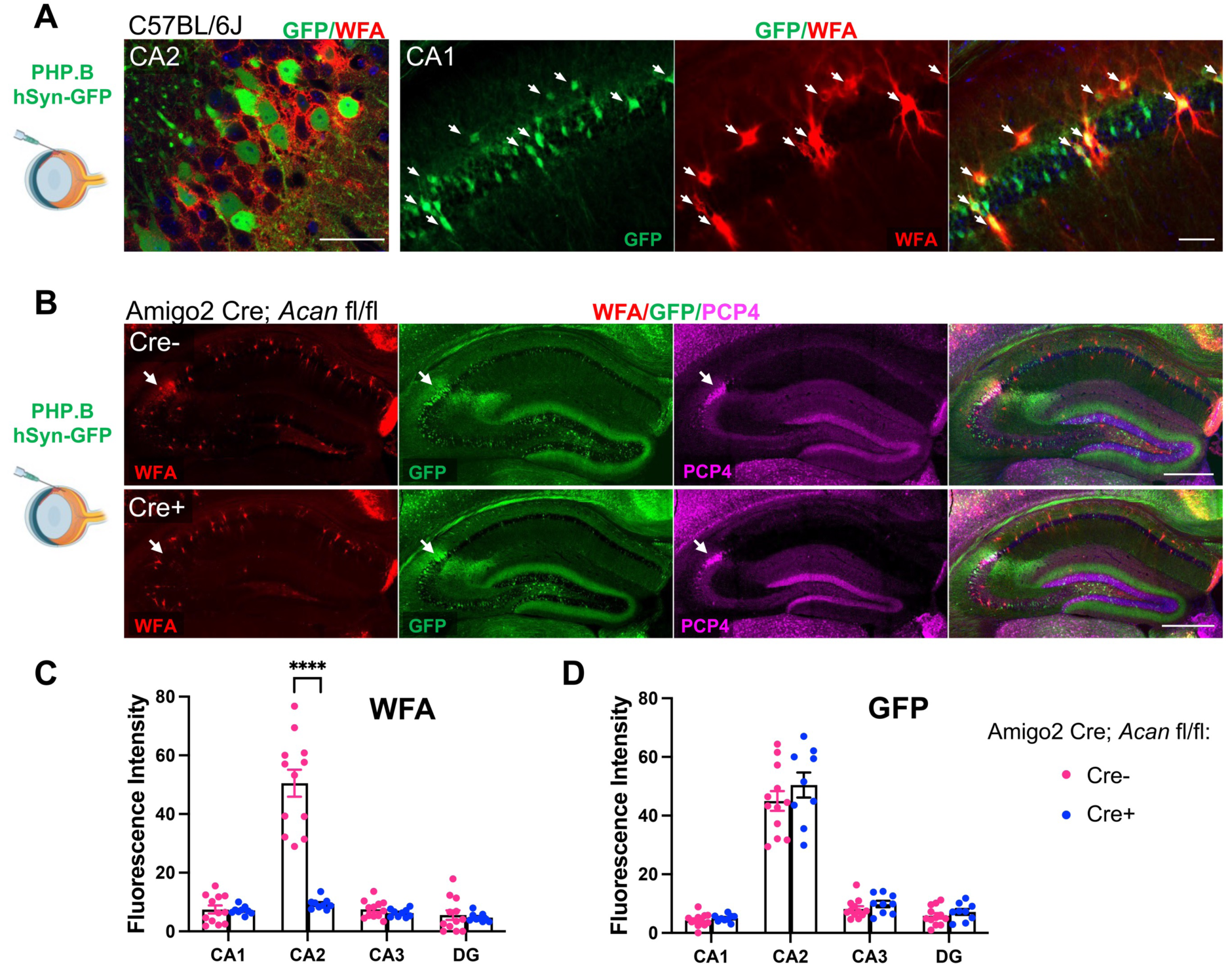
Perineuronal nets do not facilitate AAV expression in CA2. A. Following retro-orbital injection of PHP.B-hSyn-GFP, GFP expression is seen in cells that also express perineuronal nets, labeled with WFA, including in pyramidal neurons of CA2 (left) and in presumptive parvalbumin expressing interneurons in CA1 (right, colabeled cells marked by arrows). B. In Amigo2 Cre; *Acan* fl/fl animals, Acan, and thus PNNs, are deleted from CA2 neurons. Arrows in each image demark CA2, as defined by PCP4 expression. C. Fluorescence intensity of WFA stain is significantly less in CA2 of Cre+ Amigo2; *Acan* fl/fl compared to Cre-controls (subfield: F(3,76)=43.84, *p*<0.0001; genotype: F(1,76)=55.64, *p*<0.0001; interaction: F(3,76)=18.04, *p*<0.0001). D. GFP expression does not differ in animals lacking PNNs in CA2 (subfield: F(3,76)=205.0, *p*<0.0001; genotype (F1,76)=2.55, *p*=0.11; interaction: F(3,76)=0.55, *p*=0.65). Repeated-measured two-way ANOVAs with Bonferroni multiple comparisons used for all analyses, and results of multiple comparisons tests are shown on graphs. Scale bars in A = 50 μm, B=500 μm.

### Primary glycan receptors and an AAV2 co-receptor are enriched in CA2

Returning to the question of other factors that may contribute to AAV transduction, we considered glycan receptors including N-linked sialic acid and galactose, and HSPGs, which have been described as the primary cell surface receptors for several AAVs (Dhungel et al., 2020). We stained hippocampal sections from C57BL/6J mice with the lectin *Sambucus Nigra* agglutinin (SNA), which binds terminal sialic acid residues (Li et al., 2020), and found significant overlap of staining for SNA and GFP following injection with either the PHP.B or CAP-B10 variant. Quantification of SNA staining showed significantly greater expression in CA2, defined by PCP4 (data not shown), than all other hippocampal areas (Fig. 6A). As Cre+ MR fl/fl animals lack AAV transduction, we measured SNA fluorescence intensity in these mice and found that Cre+ animals had significantly less SNA expression than Cre-animals (Fig. 6B). We additionally probed AAVR KO tissue for SNA and found no difference in SNA expression between AAVR KO animals and WT littermates (Fig. 6C). We next stained tissue with lectin *Maackia amurensis* lectin I (MAL I), which detects terminal galactose (Bai et al., 2001; Li et al., 2020), and found significant enrichment on CA2 cell membranes, although staining appeared lighter than with SNA (Supp. Fig. 4A). As with SNA, MAL I staining was not enriched in CA2 of Cre+ MR fl/fl animals but appeared similar to staining in C57BL/6J animals in AAVR KOs (Supp. Fig. 4B-C).

**Figure 6.**
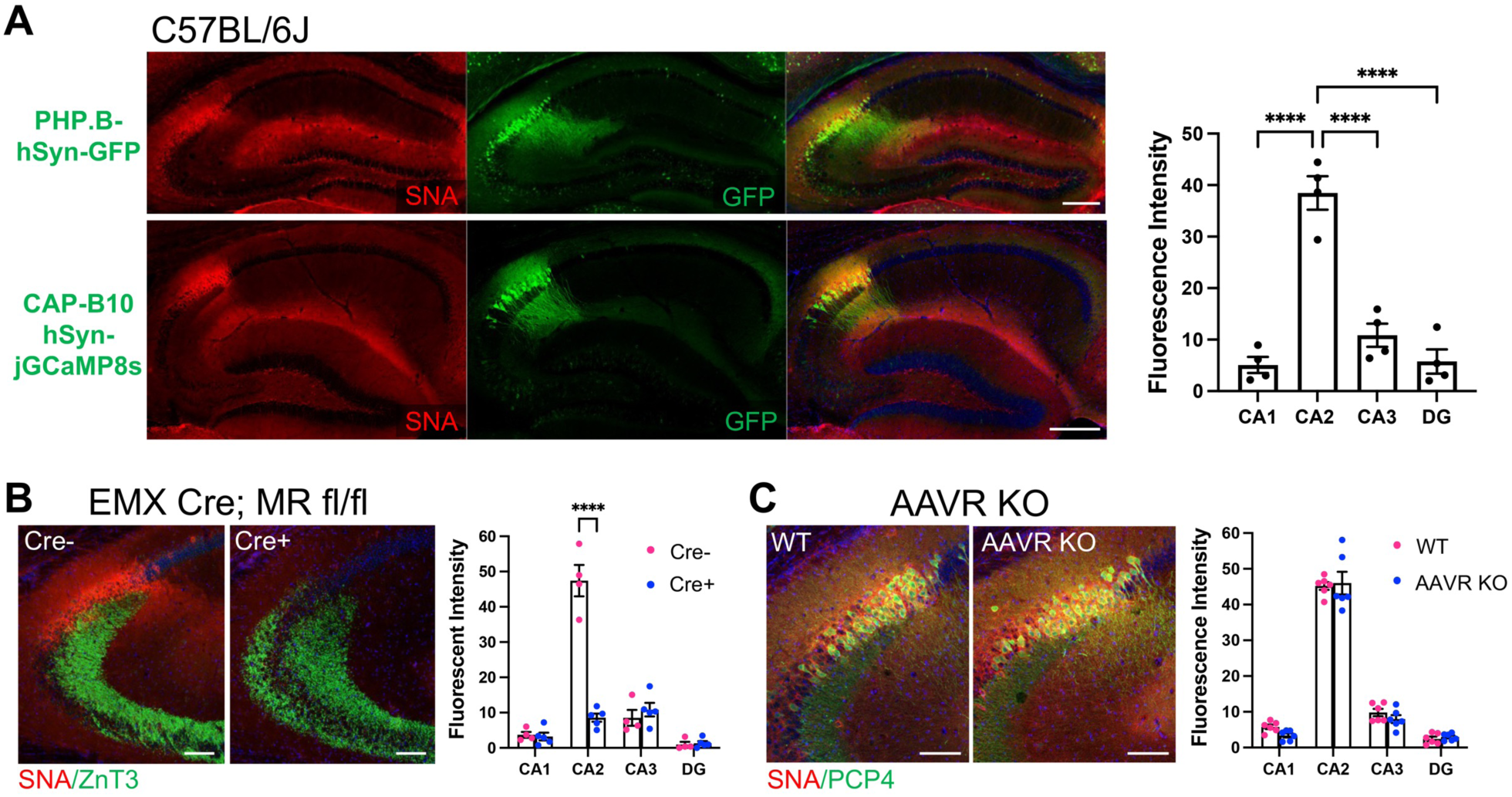
*Sambucus Nigra* agglutinin (SNA), which recognizes N-linked sialic acid glycans and acts as a primary glycan receptor for AAVs enriched in CA2. A. SNA staining colocalizes with GFP expression following both PHP.B- and CAP-B10 injection. SNA fluorescence intensity is significantly higher in CA2 than all other subfields (F(3,9)=98.79, p<0.0001, one-way ANOVA; results of Tukey’s multiple comparisons tests shown on graph). B. In Cre+ MR fl/fl animals, SNA fluorescence is decreased in CA2, defined as the pyramidal cells at the distal end of the ZnT3-expressing mossy fibers (genotype: F(1,7)=20.89, *p*=0.0026; subfield: F(3,21)=142.4, *p*<0.0001; interaction: F(3,21)=94.62, *p*<0.0001, repeated-measures two-way ANOVA with Bonferroni multiple comparisons tests, shown on graph). C Immunofluorescence for SNA does not differ between AAVR KO animals and WT littermates (genotype: F(1,10)=0.3721, *p*=0.56; subfield: F(3,30)=544.2, *p*<0.0001; interaction: F(3,30)=0.88, *p*=0.47).

**Figure 7.**
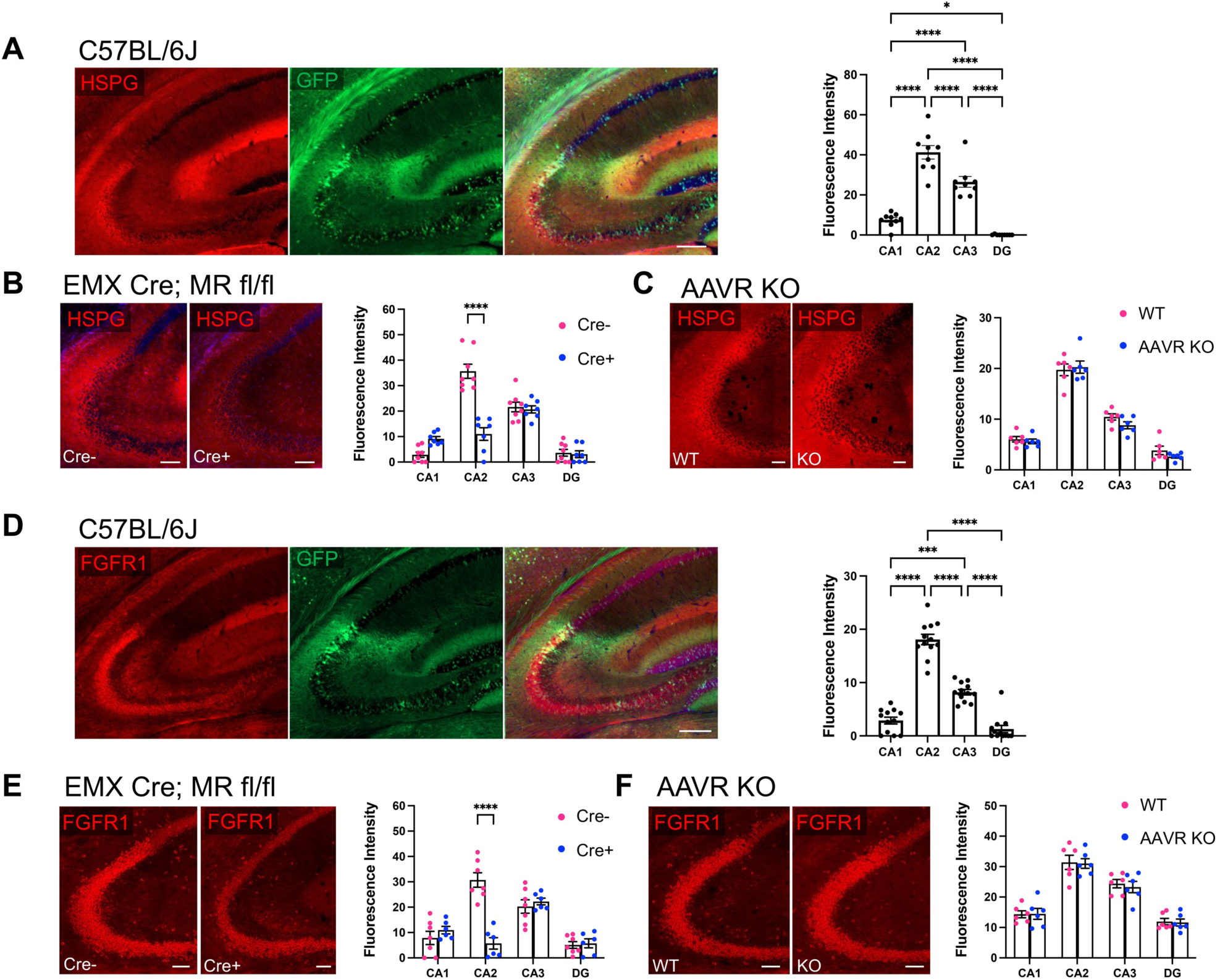
HSPG and FGFR1 stain is enriched in CA2 and colocalizes with (PHP.B) GFP. A. HSPGs, which are considered a primary glycan receptor for AAVs, showed significantly greater expression in CA2 compared with other subfield and coloalized with GFP expression following PHP.B-hSyn-GFP injection. CA3 also showed greater HSPG staining than CA1 and DG (F(3,24)=105.6, p<0.0001, RM one-way ANOVA; results of Tukey’s multiple comparisons tests shown on graph). B. HSPG staining is significantly decreased in CA2 of EMX-Cre+; MR fl/fl animals compared with Cre-animals (subfield: F(3,39)=97.47, *p*<0.0001; genotype: F(1,13)=7.77, *p*=0.015; interaction: F(3,39)=42.59, *p*<0.0001; RM two-way ANOVA, Bonferroni’s multiple comparisons results shown on graph). C. HSPG staining level was similar between AAVR KO animals and WTs (subfield: F(3,30)=199.8, *p*<0.0001; genotype: F(1,10)=1.13, *p*=0.31; interaction: F(3,30)=0.84, *p*=0.48, RM two-way ANOVA). D. Immunostaining for FGFR1, which has been described as an AAV co-receptor, in tissue from C57BL/6J animals injected with PHP.B-hSyn-GFP shows colocalization of FGFR1 and GFP. Fluorescence intensity for FGFR1 was significantly greater in CA2 than all other subfields, and CA3 showed increased expression relative to CA1 and DG (F(3,33)=166.9, *p*<0.0001, RM one-way ANOVA; results of Tukey’s multiple comparisons tests shown on graph). E. FGFR1 expression in CA2 is significantly decreased in Cre+ MR fl/fl aninmals compared to Cre-animals (subfield: F(3,33)=31.93, *p*<0.0001; genotype: F(1,11)=5.31, *p*=0.041; interaction: F(3,33)=26.44, *p*<0.0001; Bonferroni’s multiple comparisons results shown on graph). F. FGFR1 expression is similar in all subfields between AAVR KOs and WT littermates (subfield: F(3,30)=167.7, *p*<0.0001; genotype: F(1,10)=0.048, *p*=0.83; interaction: F(3,30)=0.14, *p*=0.93, RM two-way ANOVA). Scale bar in A, D=200 μm. Scale bars in B, C, E, F = 100 μm. *p<0.05, ***p<0.001, ****p<0.0001.

HSPGs also act as primary glycan receptors. We stained brain sections from PHP.B-hSyn-GFP C57BL/6J mice for HSPGs using the 10E4 antibody that recognizes heparan sulfates of HSPGs (van der Born et al., 2005) and found significant enrichment in CA2 relative to all other subfields (Fig. 7A). Of note, CA3 also showed significantly greater expression of HSPGs than CA1 and DG. Again, we probed tissue from MR fl/fl mice and AAVR KO mice. We found that Cre+ MR fl/fl mice had significantly less HSPG stain in CA2 than Cre-animals (Fig. 7B). However, there was no difference in HSPG staining between WT and AAVR KO animals (Fig. 7C). Thus, similar to the terminal sialic acid and galactose staining with SNA and MAL I, respectively, HSPGs likely represent another primary glycan receptor that is enriched in CA2 and contributes to the disproportionate tropism of CA2 neurons. However, given the level of SNA, MAL I and HSPG staining in WT and AAVR KO animals, it is likely that, although primary glycan receptor expression contributes to AAV transduction, AAVR plays a larger, if not requisite, role in AAV transduction.

Fibroblast growth factor 1 (FGFR1), has been described as an AAV co-receptor and significantly increases AAV transduction, as shown for AAV2 (Qing et al., 1999). To determine whether FGFR1 could be another factor favoring CA2 transduction, we immunostained tissue from PHP-B-hSyn-GFP-injected animals for FGFR1. In C57BL/6J mice, we found that CA2 had greater expression of FGFR1 than any other subfield, and CA3 had greater expression than CA1 and DG (Fig. 7D). As with the primary glycan receptors, we measured FGFR1 expression level in Cre- and Cre+ MR fl/fl animals as well as WT and AAVR KO animals. We again found that FGFR1 was significantly decreased in Cre+ MR fl/fl animals relative to Cre-animals (Fig. 7E), yet WT and AAVR KO animals did not differ in FGFR1 expression levels (Fig. 7F).

Finally, given the previous findings that the LY6A receptor on cerebral vasculature mediates the brain permeability of PHP-type variants following systemic injection (Huang et al., 2019), we stained hippocampal tissue for LY6A. We found no enrichment of LY6A in the vicinity of CA2 to suggest greater brain permeability of systemically-administered AAVs near CA2 (Supp. Fig. 4D).

### AAV6 effectively transduces cells with minimal AAV binding factors

So far, we have found that several AAV serotypes and variants preferentially transduce CA2 pyramidal cells, but AAV6 seems to show great specificity to the site of injection without preference for CA2. We have also found that whereas CA2 in WT animals is enriched with several factors that contribute to AAV transduction, Cre+ MR fl/fl animals do not show such enrichments. We sought to determine whether AAV6 is capable of overcoming the low levels of AAV binding and entry factors in Cre+ MR fl/fl animals. Therefore, we focally injected AAV6-hSyn-GFP into CA2 of Cre- and Cre+ MR fl/fl animals. We found that CA2 neurons could be transduced in Cre+ MR fl/fl animals, and GFP expression level did not differ from that in Cre-animals (Fig. 8). Therefore, for intrahippocampal injections, AAV6 may be a valuable tool to provide robust gene expression with limited spread outside of the injection area.

**Figure 8.**
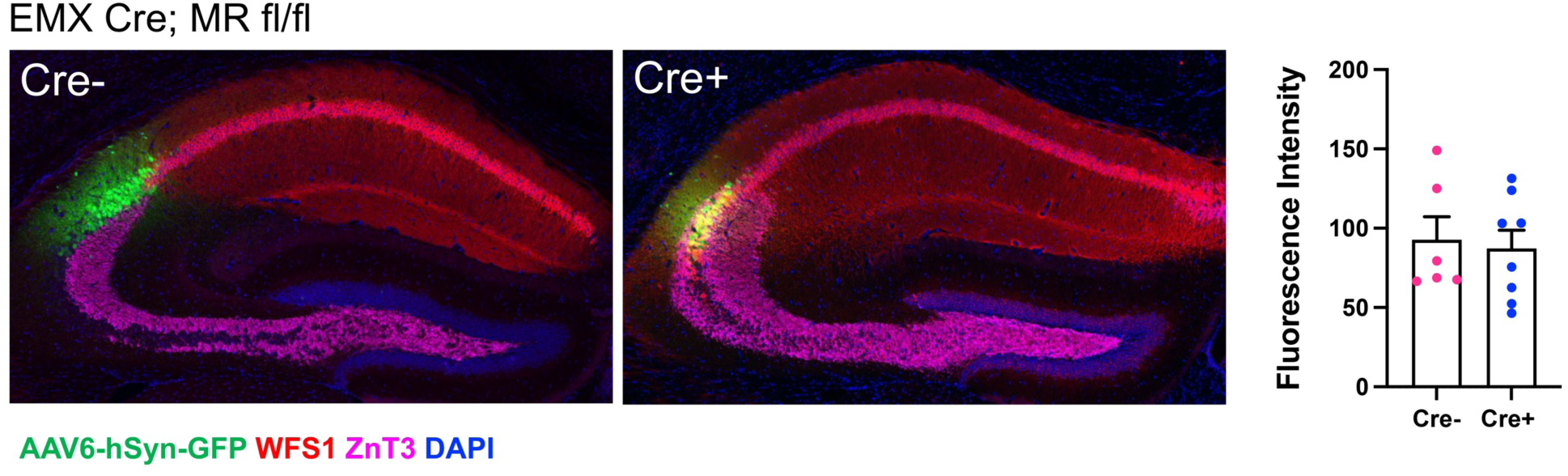
Focal injection of AAV6-hSyn-GFP into CA2 permits GFP expression in EMX Cre+; MR fl/fl animals at levels not significantly different from Cre-animals. Mossy fibers are immunopositive for ZnT3, and CA1 pyramidal cells are immunopositive for WFS1. In Cre-animals, CA2 pyramidal cells reside between the mossy fiber stain and the WFS1 stain, but in Cre+ animals, ZnT3 and WFS overlap in CA2. Fluorescence intensity of GFP in CA2 delivered by AAV6 intrahippocampal injection was not significantly different between Cre- and Cre+ animals (t(12)=0.29, *p*=0.77, two-tailed unpaired t-test).

## DISCUSSION

AAVs are among the most commonly used tools in neuroscience, allowing researchers to express effector molecules, fluorophores, recombinases, etc., in discrete populations of neurons to facilitate study of those cells (Nectow and Nestler, 2020). In experiments from our lab and from observations in the literature, we have noted that CA2 pyramidal cells seem to have a high affinity for AAVs. Using several different AAV serotypes and variants injected into hippocampus or systemically, and carrying a variety of different genetic materials, CA2 neurons seem to disproportionately express the AAV-delivered material. In this study, we tested the validity of this observation by measuring expression level of GFP when packaged in each of 9 AAV serotypes and variants delivered via either intrahippocampal or retro-orbital injection. We found that most (6 of the 9) disproportionately transduced CA2 pyramidal cells, although 2 did not and 1 produced a very little GFP expression, making it difficult to draw a conclusion. An additional retro-orbitally injected variant, CAP-B10-hSyn-jGCaMP8s, produced remarkably selective targeting of CA2 (Supp. Fig. 2). In attempts to identify why so many serotypes/variants seem to target CA2 neurons, we considered other unique properties of CA2 that could impact AAV tropism as well as known AAV binding and entry factors as possible mechanistic factors.

Pyramidal neurons in CA2 are unique among those in the other hippocampal regions in that they express PNNs, which among other functions, are thought to buffer the diffusion of ions and molecules. In addition, negatively charged chondroitin sulfate molecules make up PNNs (Morawski et al., 2015; Fawcett et al., 2022). These two factors led us to hypothesize that PNNs, that surround CA2 neurons in a spongy, negatively charged matrix, allow for increased exposure to the positively charged AAV particles. Consistent with this hypothesis, EMX Cre+ MR fl/fl animals, which lack CA2 PNNs as well as many other CA2 markers, also lacked GFP expression following PHP.B-hSyn-GFP injection. In addition, we noted that neurons in hippocampus that stained for PNNs, including both CA2 pyramidal neurons and presumptive parvalbumin-expressing interneurons, also expressed PHP.B-delivered GFP. We tested our hypothesis using Amigo2; *Acan* fl/fl animals because Cre+ animals of this strain lack Acan, and thus PNNs, but retain other CA2 markers. We injected Cre- and Cre+ mice of this strain with PHP.B-hSyn-GFP and measured GFP expression. Contrary to our hypothesis, Cre+ animals, which lack CA2 PNNs, expressed GFP in CA2 at the same level as Cre-animals, which have CA2 PNNs. Thus, at least one unique feature of CA2 neurons – expression of PNNs – is not responsible for our observations.

We next considered receptors and factors that contribute to AAV binding and internalization and asked whether any of them are enriched in CA2, starting with AAVRs (Dhungel et al., 2020). AAVRs have been described as essential for AAV transduction; deletion has been shown to render cell lines immune to infection by several AAV serotypes and impair transduction of some AAVs *in vivo* (Pillay et al., 2016; Dudek et al., 2018). Conversely, AAVR overexpression increases AAV transduction *in vitro* and *in vivo* (Pillay et al., 2016, Zengel et al., 2023). Since the discovery of the AAVR, interest has grown in identifying the interaction domain(s) between AAVR and capsids. Such domains have been identified on AAVR, and distinct domains reportedly interact with AAVs according to serotype (Pillay et al., 2017; Zhang et al., 2019; Jang et al., 2022).

We found that CA2 neurons have the highest expression level of AAVR among the hippocampal subfields, and neurons with greater immunofluorescence for AAVR also appear to have greater GFP expression following PHP.B-hSyn-GFP injections. AAVR knockout animals were devoid of GFP expression in any region following PHP.B or PHP.eB injection, and GFP expression following intrahippocampal injection of each serotype/variant was reduced by greater than 90%. Therefore, our finding corroborates previous studies stating the essential role of AAVR in transduction. However, the interaction between AAV capsids and AAVR is facilitated by other glycan and proteinaceous factors.

Cell surface glycans are thought to act as AAV attachment factors by binding specific viral capsid domains, then presenting the virion to the AAVR (Meyer and Chapman, 2022). These glycans include HSPGs, N-linked sialic acid, and N-linked galactose, and we found each of these to be enriched in CA2. HSPGs are similar to CSPGs that make up PNNs, but AAVs particularly attach to heparan sulfate containing proteoglycans, as first shown in the case of AAV2 (Summerford and Samulski, 1998). Consistent with previous findings (Fuxe et al., 1994), we found that HSPG staining is increased in CA2, paralleling the increased expression of AAV2-delivered GFP there (Fig. 1). Among the several HSPG members, glypican 1, a membrane anchored HSPG, shows increased mRNA in expression in CA2 and CA3 within hippocampus, thus supporting the enriched HSPG staining that we detected (Farris et al., 2019).

N-linked sialic acid, detectable by staining with the lectin SNA, acts as a primary glycan receptor for AAV1, 5 and 6 (Wu et al., 2006; Walters et al., 2001). We found SNA staining to be significantly higher in CA2 than any other subfield. Additionally, N-linked terminal galactose, detectable by staining with MAL I (Bai et al., 2001; Li et al., 2020), acts as the primary glycan receptor for AAV9 (Shen et al., 2011); MAL I staining was significantly higher in CA2 than surrounding subfields. Interestingly, these glycan enrichment patterns across the subfields (plotted in Fig. 6A and Supp. Fig. 4) are mimicked by the relative GFP expression following injection with the PHPs or CAP-B10 variants (plotted in Fig 2B and Supp. Fig 2, respectively), suggesting an important faciliatory role of these glycan receptors in engaging the richly expressed AAVR in CA2.

AAV2 showed the greatest tropism for CA2 among the injected AAVs in this study, likely due to enrichment of the binding factors that it uses. The primary glycan receptor for AAV2 is HSPGs (Summerford and Samulski, 1998), but AAV2 additionally makes use of FGFR1 as a proteinaceous co-receptor. Expression of both factors are required for efficient transduction (Qing et al., 1999). As with many others serotypes, AAV2 also requires AAVR (Pillay et al., 2016). We found that all of the factors mentioned here (HSPG, FGFR1, AAVR) were expressed significantly higher in CA2 than surrounding subfields, such that CA2 is primed for expression of AAV2-delivered genetic material. The significance of this affinity for viruses remains unknown.

By contrast, AAV6 showed the lowest tropism for CA2. AAV6 is thought to use HSPGs and N-linked sialic acid as primary glycan receptors. Blockade or removal of either of these receptors does not significantly affect transduction, and the structure of AAV6 shows multiple glycan binding sites (Wu et al., 2006; Mietzsch et al., 2014), suggesting that AAV6 is a more promiscuous AAV. This serotype reportedly makes use of the epidermal growth factor receptor (EGFR) as a co-receptor, but expression of EGFR appears universally low in hippocampus (Farris et al., 2019). We found that AAV6 requires AAVR for effective transduction, but in its complete absence in the AAVR KO animals, AAV6 produced significantly greater GFP expression than other serotypes, suggesting that AAV6 may use a variety of receptors. GPR108 is another reported AAV6 entry factor, but expression is uniformly low expression throughout hippocampus (Dudek et al., 2020; Farris et al., 2019). Little else of the binding and entry factors used by AAV6 are known, but perhaps the multiple glycan binding sites provide a clue that AAV6 can capitalize on low expression levels of binding and entry factors to nonetheless transduce cells. In support of this, CA2 neurons in EMX Cre+; MR fl/fl mice, which have low expression levels of AAVR, HSPGs and glycan receptors, were effectively transduced by focal injection of AAV6-hSyn-GFP into CA2.

The PHP variants were engineered by directed evolution from AAV9 to enhance blood-brain permeability and increase neuronal tropism. Images from these studies hinted at tropism toward CA2 (Chan et al., 2017; Goertsen et al., 2022), which was later validated when PHP.eB variant was injected retro-orbitally or into the lateral ventricle to reveal selective CA2 expression (Okamoto et al., 2023). The CAP-B10 variant was evolved from PHP.eB to further enhance neuronal tropism and minimize peripheral organ tropism (Goertsen et al., 2022). Following retro-orbital injection of PHP.B, PHP.eB and CAP-B10 variants, we found significantly greater transduction of neurons in CA2 than other hippocampal subfields. As these variants are derived from AAV9, they may use the same primary glycan receptor, the terminal galactose glycan (Jang et al., 2022), which we found to be expressed at higher levels in CA2.

The CAP-B10 variant appears to have produced the most selective transduction of CA2 neurons, supporting the use of this variant for non-invasive administration of CA2-directed genetic material. In a previous study, CA2-specific promoters were used to direct expression of AAVs in CA2 but still required intrahippocampal injection (Peng et al., 2023). Given the curved shape of hippocampus and small size of CA2 in mice, a systemically available AAV that can selectively target CA2 neurons would greatly facilitate CA2 study. Lateral ventricle delivery of the PHP.eB variant to target CA2 greatly advanced progress toward that goal (Okamoto et al., 2023). CAP-B10 may be the next step in refining CA2 targeting in a non-invasive manner. In addition, because CAP-B10 is capable of crossing the blood-brain barrier in non-human primates (Goertsen et al., 2022), this variant may provide an attractive tool to potentially target CA2 neurons for gene therapy.

## Acknowledgements

We thank the expert staff at both the NIEHS Fluorescence Microscopy and Imaging Center, the NIEHS animal care staff for all their support, Sebastian Swiggum for technical assistance.

---

Illustrations created with BioRender.com

**Supplemental Figure 1.**
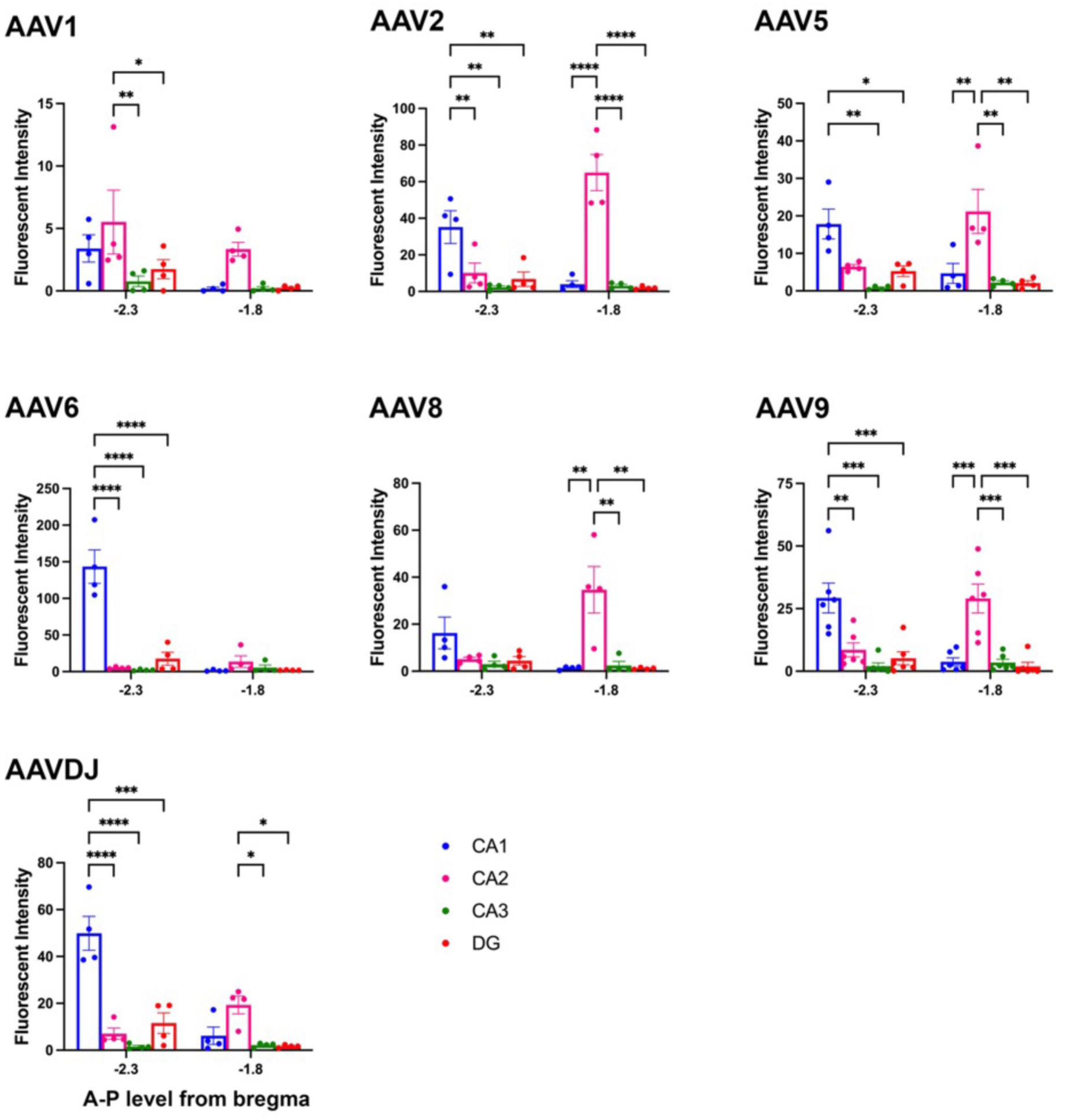
Intrahippocampally injected AAVs showed differential preference for CA2 expression according to serotype. GFP fluorescence intensity was measured in each hippocampal subfield at each of the injection site (−2.3 mm AP) and a site in dorsal hippocampus (−1.8 mm AP). Image acquisition settings were held constant within each serotype but differed across serotypes. Fluorescence intensity values were compared using two-way ANOVAs with Sidak’s multiple comparisons tests. Main effects and interactions are as follows and results of Sidak’s multiple comparison’s tests are shown on each graph. AAV1: main effect of AP level: F(1,3)=5.40, *p*=0.10, main effect of subfield: F(3,9)=5.38, *p*=0.021, interaction: F(3,9)=1.10, *p*=0.40. AAV2: AP level: F(1,3)=0.99, *p*=0.39, subfield: F(3,9)=14.34, *p*=0.0009, interaction: F(3,9)=42.28, *p*<0.0001. AAV5: AP level: F(1,3)=0.0009, *p*=0.98, subfield: F(3,9)=9.39, *p*=0.0039, interaction: F(3,9)=11.44, *p*=0.0020. AAV6: AP level: F(1,3)=33.18, *p*=0.010, subfield: F(3,9)=22.59, *p*=0.0002, interaction: F(3,9)=29.27, *p*<0.0001. AAV8: AP level: F(1,3)=0.51, *p*=0.52, subfield: F(3,9)=5.38, *p*=0.021, interaction: F(3,9)=11.78, *p*=0.0018. AAV9: AP level: F(1,5)=0.61, *p*=0.47, subfield: F(3,15)=13.76, *p*=0.0001; interaction: F(3,15)=18.26, *p*<0.0001. AAVDJ: AP level: F(1,3)=6.92, *p*=0.078, subfield: F(3,9)=24.03, *p*=0.0001, interaction: F(3,9)=22.89, *p*=0.0002.

**Supplemental Figure 2.**
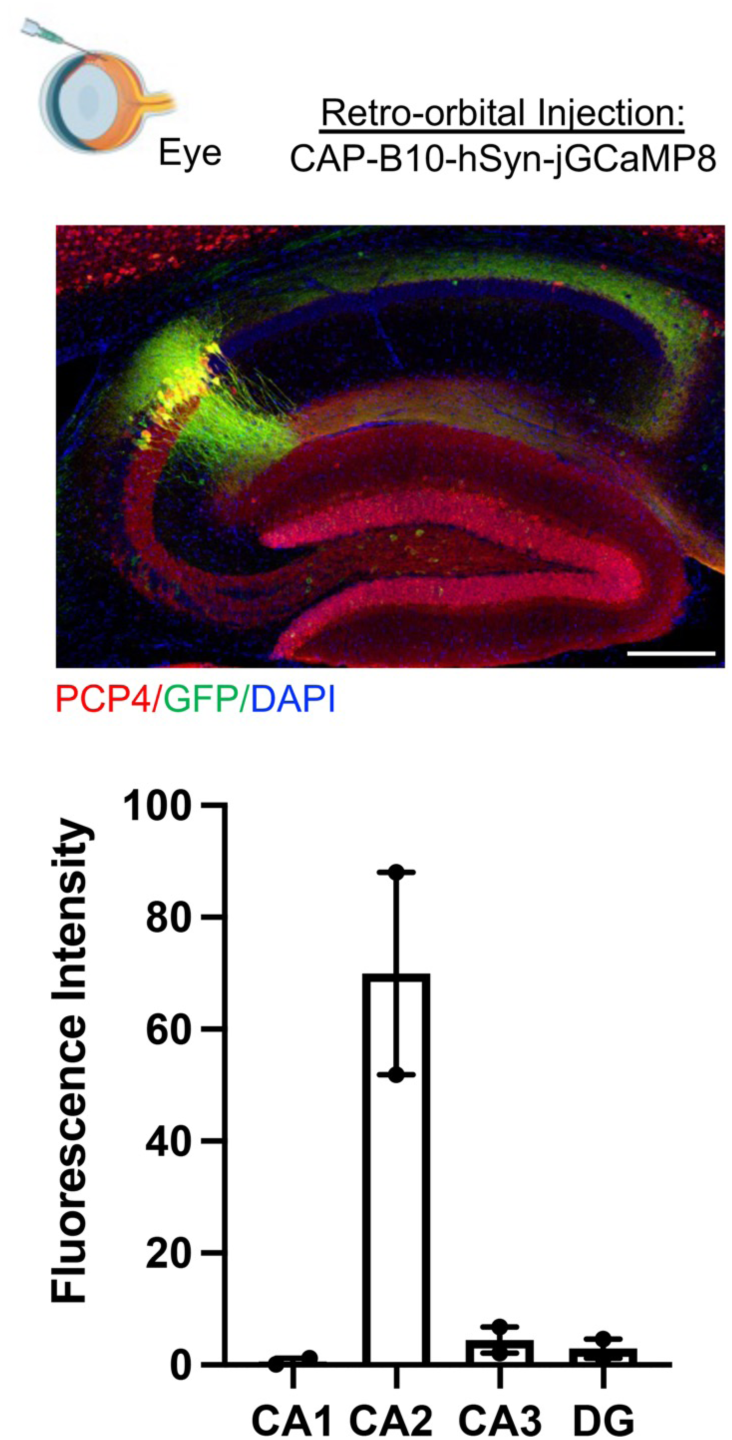
CAP-B10-hSyn-jGCaMP8s shows prominent tropism for CA2 neurons. Statistics are not included due to the small number of animals (N=2 mice).

**Supplemental Figure 3.**
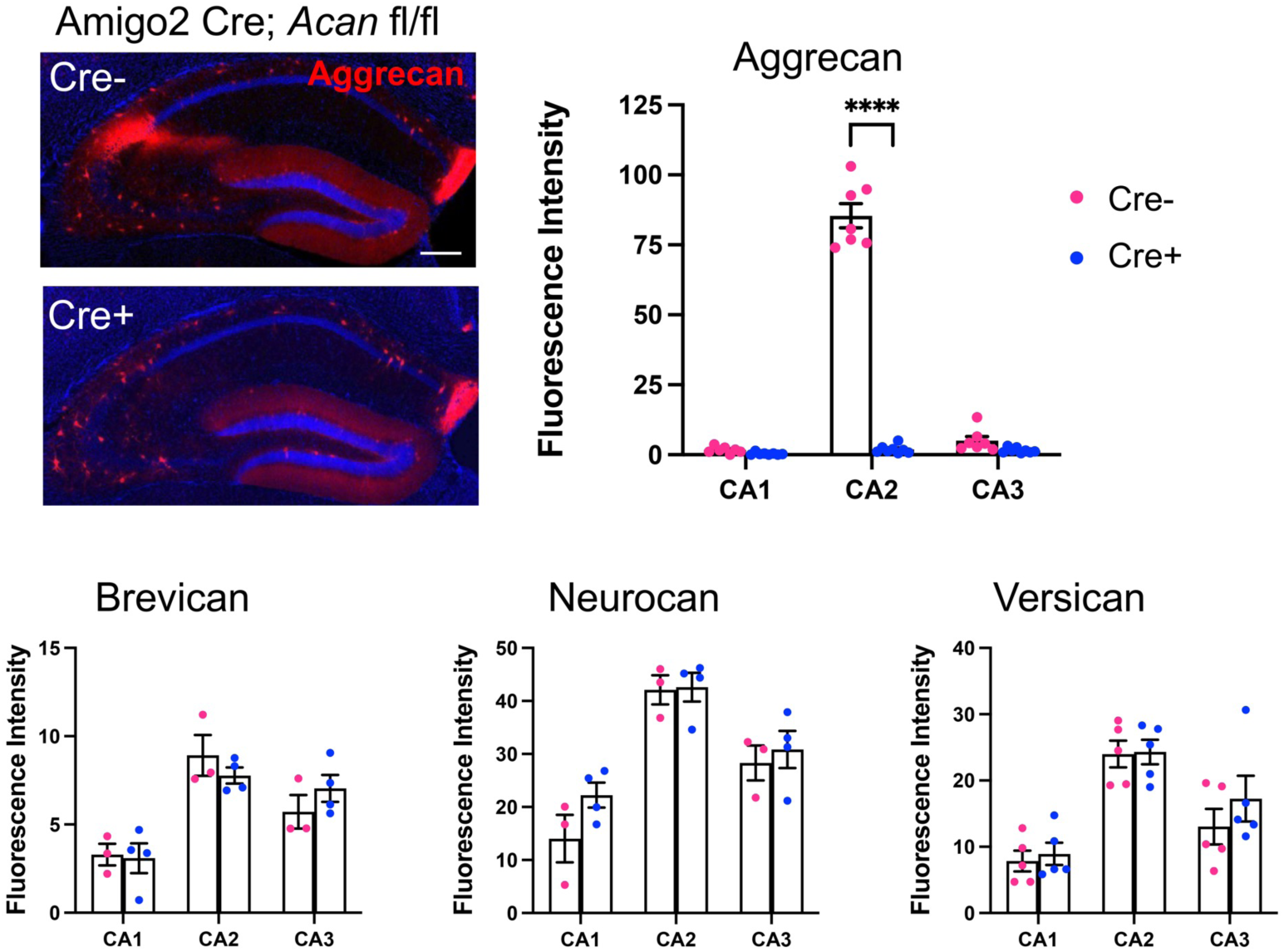
Aggrecan expression is significantly decreased in Amigo2 Cre+ *Acan* fl/fl animals. Expression on brevican, neurocan and versican are not affected by *Acan* deletion in CA2 (Aggrecan: subfield: F(2,26)=491.7, *p*<0.0001; genotype: F(1,13)=283.1, *p*<0.0001; interaction: F(2,26)=468.2, *p*<0.0001; Brevican: subfield: F(2,10)=82.21, *p*<0.0001; genotype: F(1,5)=4.9×10^-5^, *p*=0.9948; interaction: F(2,10)=4.70, *p*=0.036; Neurocan: subfield: F(2,10)=46.04, *p*<0.0001; genotype: F(1,5)=1.15, *p*=0.33; interaction: F(2,10)=1.25, *p*=0.33; Versican: subfield: F(2,16)=36.91, *p*<0.0001; genotype: F(1,8)=0.58, *p*=0.47; interaction: F(2,16)=0.64, *p*=0.54. Scale bar = 250 μm.

**Supplemental Figure 4.**
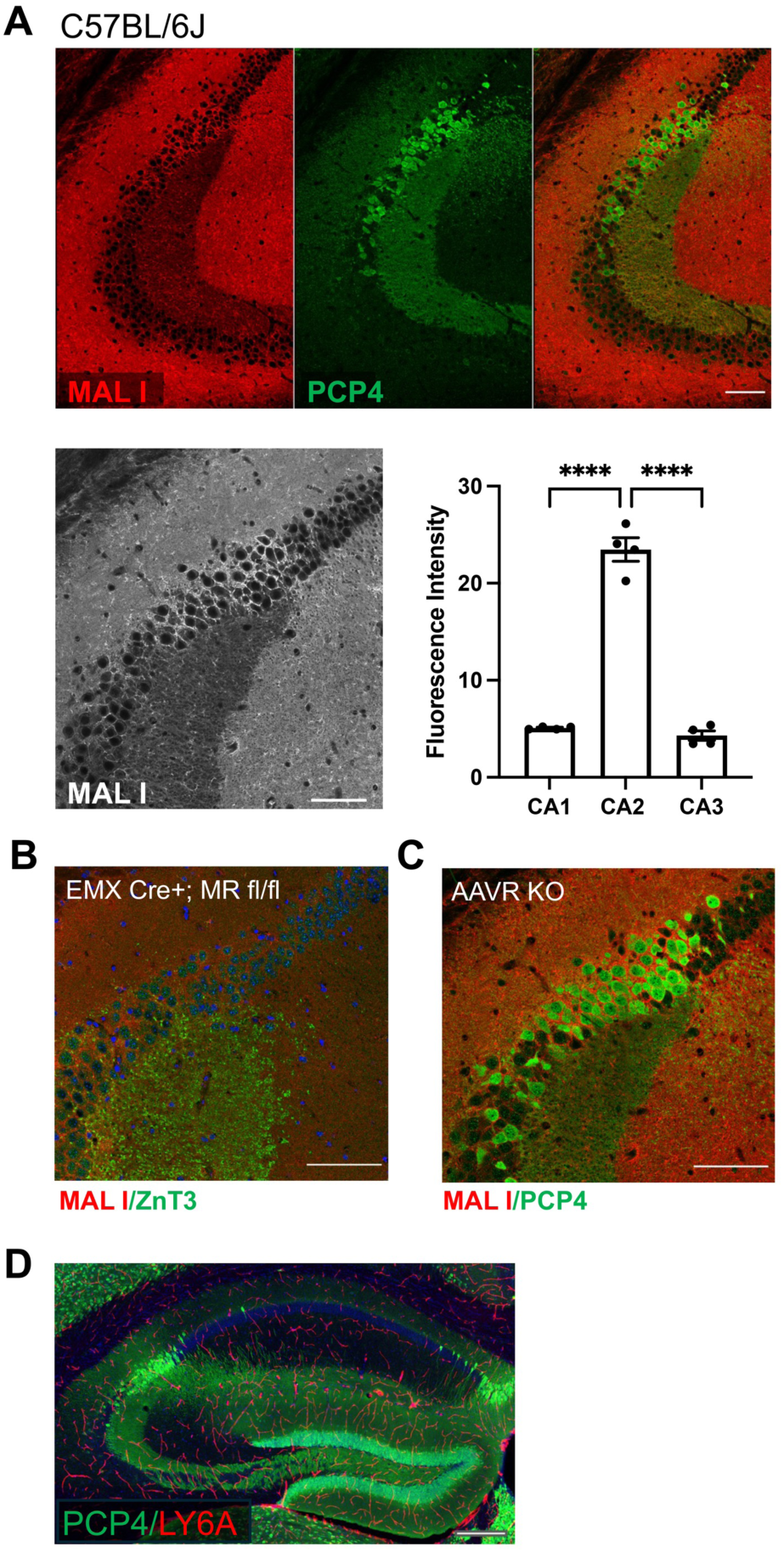
A. MAL I, which recognizes terminal galactose glycans shows light stain on the cell membrane of CA2 pyramidal cells, encircling PCP4-positive cells Grayscale image below shows more clearly than color used for colocalization. MAL I staining was significantly higher in CA2 than either CA1 or CA3 (F(2,6)=171.8, p<0.0001; one-way ANOVA). B-C. MAL I staining appeared decreased in Cre+ MR fl/fl animals (B) but appeared similar to that in C57BL/6J animals (C). D. The PHP receptor, LY6A, on vasculature does not appear to be enriched near CA2. Scale bars = 100 μm in A, 250 μm in B.

Supplemental Video 1. Expression of GFP in an Amigo2 CreERT2; ROSA tdTomato mouse following retro-orbital injection of PHP.B-hSyn-GFP. CA2 pyramidal cells and their projections are shown in red.

Supplemental Video 2. Expression of GFP (pseudocolored as “fire”) in an Amigo2 Cre+; *Acan* fl/fl animal.

